# Diversity of culturable actinobacteria producing protease inhibitors isolated from the intertidal zones of Maharashtra, India

**DOI:** 10.1101/2020.02.14.949438

**Authors:** Neha Shintre, Ulfat Baig, Anagha Pund, Rajashree Patwardhan, Vaijayanti Tamhane, Neelima Deshpande

## Abstract

Phylogenetic diversity of culturable actinobacteria isolated from the intertidal regions of west coast of Maharashtra, India was studied using 16S rRNA gene sequencing. Total of 140 actinobacterial isolates were obtained, which belonged to 14 genera, 10 families and 65 putative species with *Streptomyces* being the most dominant (63%) genus followed by *Nocardiopsis* and *Micromonospora*. They were screened for production of extracellular protease inhibitors (PI) against three pure proteases viz. chymotrypsin, trypsin, subtilisin and one crude extracellular protease from *Pseudomonas aeruginosa*. Eighty percent of the isolates showed PI activity against at least one of the four proteases, majority of them belonged to genus *Streptomyes*. Actinobacterial diversity from two sites Ade (17°52′ N, 73°04′ E) and Harnai (17°48′ N, 73°05′ E) with varying degree of anthropological pressure showed that more putative species diversity was obtained from site with lower human intervention i.e Ade (Shannon’s H 3.45) than from Harnai (Shannon’s H 2.83), a site with more human intervention. Further, in Ade percentage of isolates not showing PI activity against any of the proteases was close to 21% and that in Harnai was close to 9%. Considering human activities in the coastal region might be contributing to increasing the organic load and in turn increasing the presence of extracellular enzymes in the intertidal environments it would be interesting to look at the association of PI production and organic load in these habitats.

## INTRODUCTION

Proteases are enzymes that catalyse proteolytic reactions and are involved in variety of biological processes like digestion, cell signalling or tumour formation (Groll et al., 2002; Koblinski et al., 2000; Overall and Blobel, 2007; Roberts et al., 2012). They are ubiquitously present across different plant, animal and microbial taxa. Two of the many roles of these enzymes in bacteria is to aid in bacterial pathogenesis (Ingmer and Brøndsted, 2009; Maeda, 1996) and bacterial predation (Martin, 2002) where they secrete hydrolytic enzymes in order to degrade prey species in the vicinity (Martin, 2002). Protease inhibitors bind reversibly or irreversibly with the enzymes, rendering them inactive. Hence, protease inhibitors (PIs) play a crucial role as defence molecules in intracellular or extracellular environment. They are classified based on either the type of protease they inhibit (e.g. serine protease inhibitor, aspartic protease inhibitor) or based on the mechanism of action (e.g. reversible or irreversible enzyme inhibitors). Various plants, animals and microorganisms produce PIs in order to deter pests and predators or secrete them as toxins for self-defence (Habib and Fazili, 2007; Hartl et al., 2011; Mourão and Schwartz, 2013). Also, PIs have high commercial potential since they are used as drugs against viral, bacterial and other parasitic infections (Karthik et al., 2014; Liu et al., 2012; Sreedharan and Bhaskara Rao, 2017; Umezawa, 1976).

Marine intercostal community harbours a variety of sedentary invertebrates and microorganisms. These organisms rely primarily on chemicals for self-defence (Engel et al., 2002; Pawlik, 1993). Additionally, as stated by Hay *et al*. the “biggest challenge for marine organisms is to obtain lunch without becoming lunch” (Hay, 2009). Protease inhibitors can play a major role in these situations since they can inhibit the action of digestive proteases present in the surroundings. Literature reports suggest that PIs are indeed produced by marine intercostal communities (Covaleda et al., 2012; Karthik and Kirthi, 2015; Karthik et al., 2014; Mourão and Schwartz, 2013) considering these facts, it might be possible to detect PIs synthesised by marine microorganisms for their defence.

Intertidal zones are exposed to air at low tides and are submerged at high tides. Thus they experience varying dry and wet periods, fluctuating temperatures and atmospheric pressure. Intertidal life forms adapt to these constantly changing environment by developing unique features. Their adaptations for survival in dynamic environments might also increase the chances of finding unique set of metabolites from them.. Moreover, it is expected that marine microorganisms will have different set of defence molecules than their terrestrial counterparts (Mourão and Schwartz, 2013; Xie et al., 2018).

Actinobacteria is one of the largest phyla in domain bacteria. These are gram positive organisms with high guanine to cytosine ratio (Trujillo, 2016). They at large are known to produce commercially important diverse metabolites. Some reports show that the marine environment has become a prime source for discovery of novel actinobacteria and novel natural products produced by them (Maldonado et al., 2005; Ward and Bora, 2006). There are few recent reports of PI production by marine actinobacteria (Karthik et al., 2014; Sreedharan and Bhaskara Rao, 2017; Sun et al., 2014) yet, to our knowledge, there are limited reports on PIs from west coast of India. Thus, in the current study, marine actinobacteria from intertidal zone were screened for presence of PIs.

It has been reported that sponges and bacteria are two main sources of enzyme inhibitors from marine environments (Ruocco et al., 2017). Some reports also suggest that many a times, metabolites obtained from sponges are actually produced by the bacterial symbionts (Haygood et al., 1999; Mehbub et al., 2014). Therefore, in the current study sponge, sediment and water samples were collected from the coast of Maharashtra and Goa for isolation of actinobacteria. Obtained isolates were subjected to molecular identification using 16S rRNA gene sequencing and isolates were screened for presence of protease inhibitors. Data was analysed to check correlations between inhibitor molecules and environmental parameters and their probable role as defence molecules.

## MATERIALS AND METHODS

### Sampling

Around 60 samples comprising of sponge, sediment and sea water were collected from the intertidal rock pools from seven different locations along the coast of Ratnagiri district from Velas, Kelshi, Ade, Anjarle, Harnai, Murud and Aare Ware. (17°57′04.9″N 73°01′43.0″E to 17°04′35.9″N 73°17′17.5″E). From Ade and Harnai, sampling was carried out in different seasons viz. pre monsoon (Mar, Apr, May), Monsoon (Early Sep), post-monsoon (Oct) and winter season (Dec, Jan, Feb). Whereas, a single sampling was done from rest of the sites. Sub-tidal sampling of sediment was also done in a single event at a place off shore to Goa (15°21′08.4″N 73°46′41.8″E). The depths of collection for this sampling were 9m, 11m, 12m and 16m. (Figure 1).

**Fig 1:**
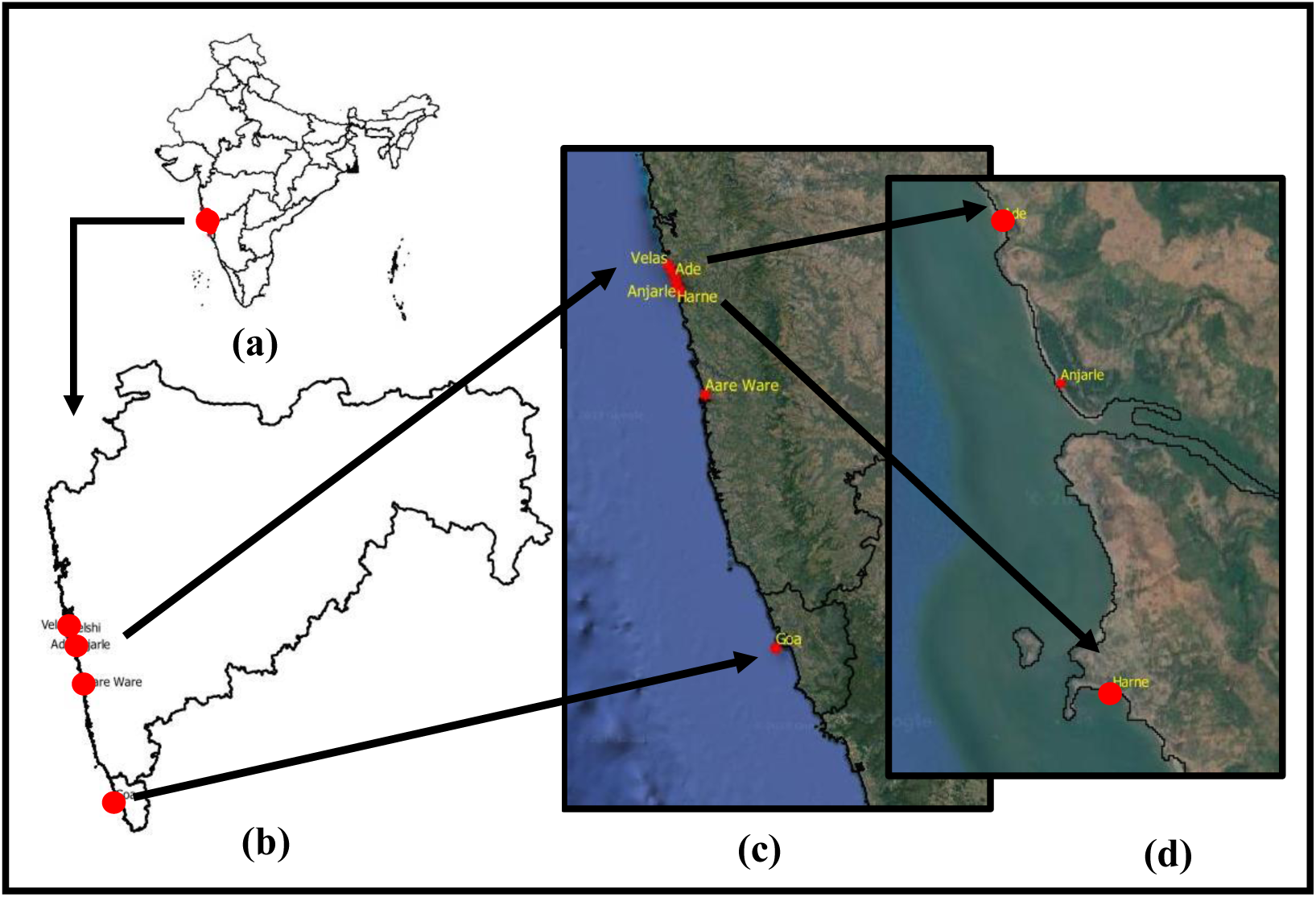
Map of west coast of India (A) illustrating sampling sites along the coast of Maharashtra and Goa (B). Corresponding positions are shown in the inset (C & D). The maps were generated using QGIS software (http://qgis.osgeo.org) and Google maps.

Small tissue samples of sponge were collected without damaging the sponge colonies or their habitat. Samples were rinsed in sterile Poor Ravan Saline (PRS) broth to remove loosely bound particles and debris and were stored in sterile collection tubes containing the same medium. Sediment and water samples in the vicinity of sponge were collected in sterile collection tubes. All the samples were transported to the laboratories in ice box and were processed fresh for microbial isolations.

### Sample Processing, Selective Isolation and Culture Maintenance

All the samples were given heat treatment at 60°C for 15 mins to reduce the load of non-sporulating bacteria. Sponge samples were homogenised in Poor Ravan Saline (PRS) medium (Watve et al., 2000) and diluted serially with 10 fold dilutions up to 10^−5^. Sediment samples were vigorously shaken for two minutes and diluted up to 10^−5^ dilutions. Sea water samples were also diluted up to 10^−5^ dilutions. 0.1 ml of undiluted, 10^−3^ and 10^−5^ dilutions were spread plated in triplicates on different growth media and incubated at room temperature for up to 21 days.

Four different growth media Sponge agar (1% macerated sponge colonies collected from the site, 50% sea water and 2.5% agar), Sea Water Agar (50% sea water and 2.5% agar), ZOBELL Marine Agar (ZOBELL, 1941) and Modified poor ravan medium (Watve et al., 2000)) were used to obtain maximum culture dependent diversity of actinobacteria. Nalidixic acid and Cyclohexamide were added to the culture medium in 25 µg ml^-1^ concentration to inhibit the growth of Gram-negative bacteria and fungi (Magarvey et al., 2004). Plates were incubated at 30°C and monitored daily for 21 days. Isolated colonies showing resemblance to typical actinobacterial colony morphology were picked up and subcultured several times for obtaining pure cultures. All the pure cultures were later stored on modified (1:4 diluted) ZOBELL marine agar at 4°C for further use.

### DNA Sequencing and Diversity Analysis

16S rRNA sequences from previous study of (Baig et al., 2020) were used to make maximum likelihood phylogenetic tree of actinobacteria used in the study. Maximum likelihood tree was constructed in IQtree (Nguyen et al., 2015). Best nucleotide substitution model was determined in model finder (Kalyaanamoorthy et al., 2017). Note support was examined using bootstrap values of 1000 iterations and *Bacillus sp*. were used as out-group. Shannon′s diversity index to measure species diversity of actinobacteria was calculated using Past (version 4.0) (Hammer, 2001).

### Screening for Protease Inhibitors (PI) using spot assay

#### Sample preparation

Detection of extracellular PIs from cell free supernatant was done as follows. Pure cultures of actinobacteria were inoculated (single colony in 150 ml broth) in ZOBELL Marine broth and incubated for 5 days at 30°C on incubator shaker with 100 rpm speed. After incubation, the cultures were centrifuged at 4000 rpm for 20 mins and cell free supernatants (CFS) were collected. 1 ml of each of these CFSs were kept in hot water bath of 70°C for 15 mins for denaturation of proteases produced by the bacteria. Heat treated CFSs were used for detection of PIs. Unprocessed X-ray films coated with gelatine were used to detect PIs. (Gelatine is degraded by various proteolytic enzymes, so, upon action of the proteases, clear zones are observed on unprocessed x-ray films at the point of contact of enzymes. On the other hand, if the gelatine layer remains intact, no clearance is observed. PIs inactivate proteases and thus, presence of protease inhibitors is marked by no clearance on the x-ray films as shown in Figure 3).

**Fig 2:**
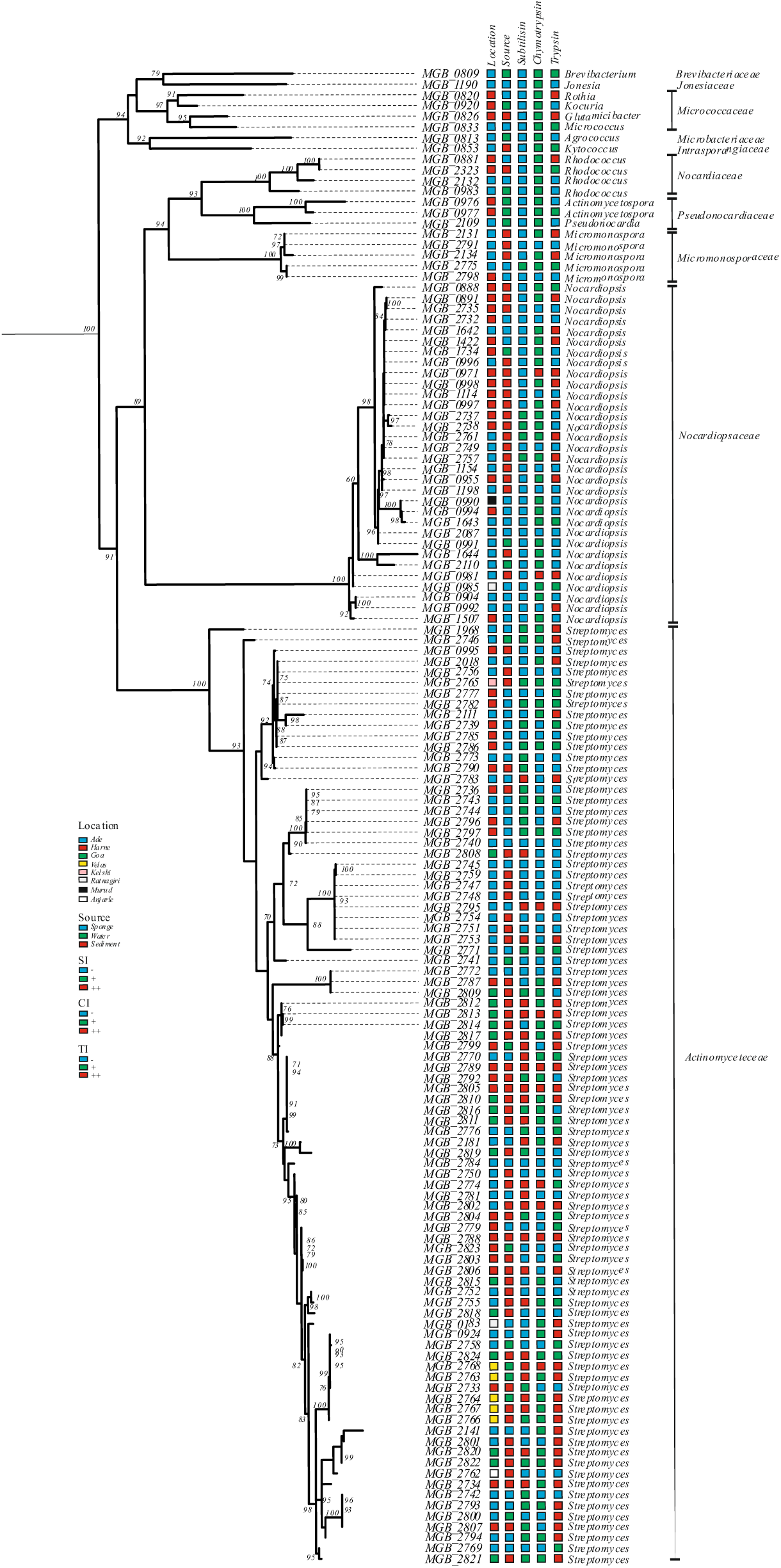
Phylogenetic relatedness of isolates in the study along with location, source of isolation and PI Profile against different enzymes. Given is a maximum likelihood tree of actinobacteria used for screening of PIs. Outgroup taxa (*Bacillus* sp.) not shown. Percent bootstrap values of 1000 iterations are provided along the nodes. The figure also depicts location and source of isolation of the isolates along with their ability for production of PIs against subtilisin (SI), chymotrypsin (CI) and trypsin (TI) where (- : negative, +: positive and ++ : strong positive)

**Fig 3:**
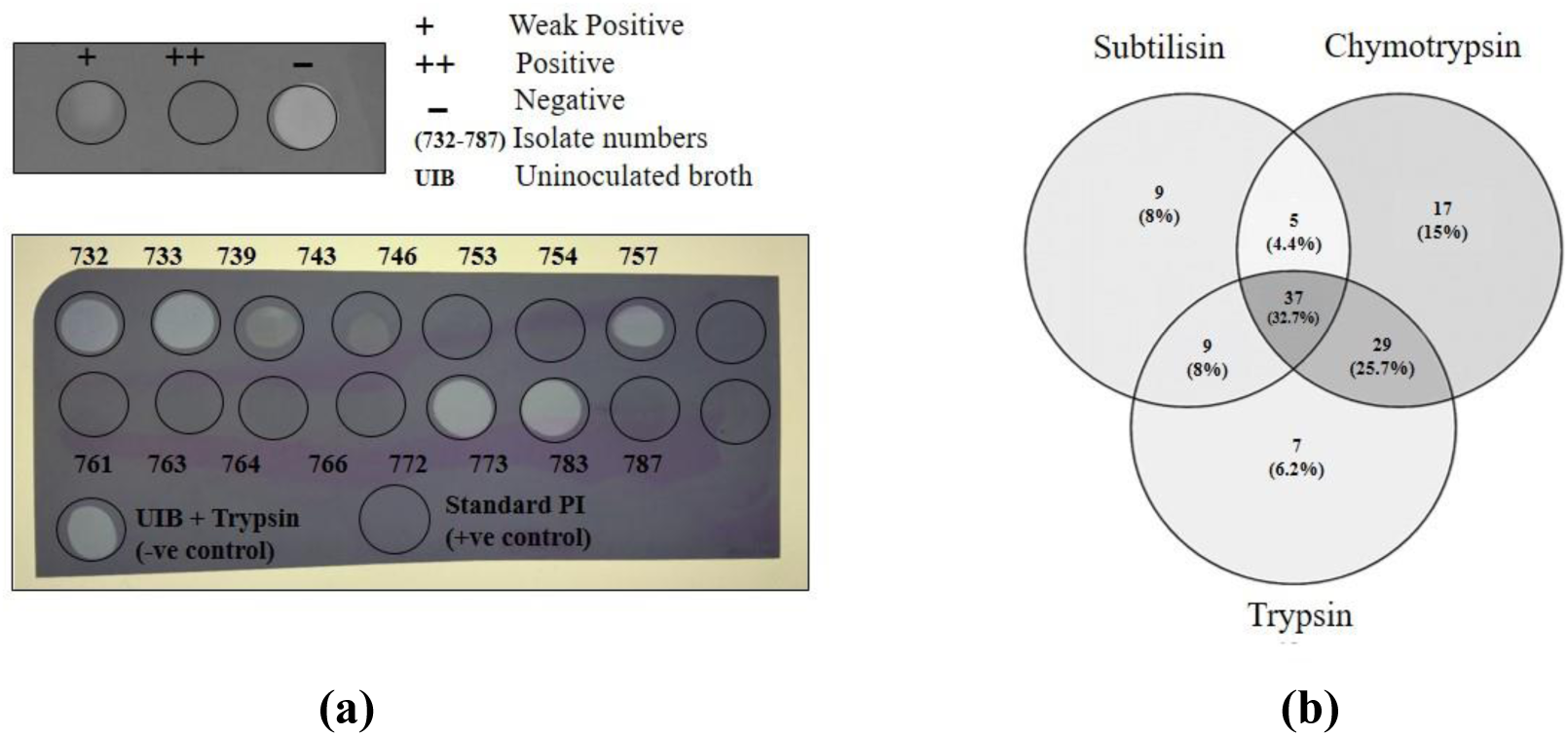
Spot assay representing PI activity against trypsin (A); and Venn diagram showing overlap of number of isolates producing subtilisin, chymotrypsin and trypsin inhibitors (B).

Proteases used for the assay included Trypsin (SRL Pvt. Ltd, Cat no.-60484) and α-Chymotrypsin (SRL Pvt. Ltd, Cat no.-35085) of bovine origin and Subtilisin (Sigma-Aldrich, Cat no.-P5380) and crude protease obtained in the laboratory from cell free supernatant of *Pseudomonas aeruginosa* (NCBI Accession number - MN044759) of bacterial origin were used to detect PIs. Amongst these enzymes, trypsin and chymotrypsin were of bovine origin. Additionally, Protease inhibitor cocktail (Sigma-Aldrich, Cat no.-P5380) was used as a positive control and un-inoculated culture broth incubated with the enzyme was used as a negative control.

#### Spot assay

Standardization of enzyme concentration was done as described by Tripathi et al., (2011).Various dilutions of pure enzyme were spotted on gelatine coated X-ray films. Lowest dilution that gave complete clearance, in turn indicating complete digestion of gelatine, was used in the assay.

Spot assay as described by (Cheung et al., 1991) for detection of PIs was carried out as follows. 10 µl of pure enzyme (100 µg ml^-1^) was incubated with 10 µl of heat treated CFS for 10 mins and then transferred to untreated X-ray-Fuji Medical X-ray, HRU grade-films. The assembly was kept undisturbed for 15 mins at room temperature to allow the enzyme substrate reaction to take place. Films were washed with running tap water and allowed to dry before recording the results.

## RESULTS

### Sample Collection

A single sampling of sediment and water was carried out from intertidal and sub-tidal regions of Aare Ware and Goa respectively. Similarly, one time sampling was carried out from intertidal zones of Velas, Kelshi, Anjarle and Murud however, at the time of sampling, sponge colonies could not be located on any of these sites and thus only sediment and water samples were collected. Whereas, at Harnai and Ade four to five distinct morphotypes of sponges were recorded. Thus, sponge sediment and water samples were collected from these sites. These sites were also selected for sampling in various seasons throughout the year. Harnai, one of the busy ports on the western coast is used for various activities like, auction of fishes on a daily basis, recreational space for the villagers or tourist hang out spot. Certain parts of intertidal rock patch in the vicinity are used as open defecation sites by the villagers and few other parts of the rock patch are used for clamp collection and fishing from the rock pools. Conversely Ade, is hardly used for any of the above purposes except for occasional fishing from the rock pools. Thus, intertidal regions of the above sites faced varying degree of anthropological disturbance with Harnai being the most and Ade being the least used site with respect to human activities.

### Distribution, Identification and Phylogeny of Actinobacteria from Sponge, Sediment and Sea Water Samples

We obtained 140 actinobacterial isolates from sponge, sediment and sea water samples. They showed high phylogenetic diversity wherein there were 65 putative species belonging to 14 genera of 10 different families. Most abundant genus amongst the isolates was *Streptomyces sp*. (∼63%) followed by *Nocardiopsis sp*. (∼22%) and *Micromonospora sp*. (∼4%). Approximately 11% isolates belonged to other genera (Table 1).

**Table 1:** Counts of putative species obtained from sponge, sediment and sea water samples.

| Family | Genus | Putative species identity | Isolates |
| --- | --- | --- | --- |
| Actinomycetaceae | Streptomyces | *Streptomyces albidoflavus* | 6 |
|  |  | *Streptomyces albogriseolus* | 8 |
|  |  | *Streptomyces atrovirens* | 1 |
|  |  | *Streptomyces aurantiogriseus* | 1 |
|  |  | *Streptomyces aureofaciens* | 1 |
|  |  | *Streptomyces cellulosae* | 1 |
|  |  | *Streptomyces champavatii* | 1 |
|  |  | *Streptomyces coeruleofuscus* | 1 |
|  |  | *Streptomyces collinus* | 1 |
|  |  | *Streptomyces diastaticus* | 3 |
|  |  | *Streptomyces euryhalinus* | 4 |
|  |  | *Streptomyces fradiae* | 6 |
|  |  | *Streptomyces geysiriensis* | 1 |
|  |  | *Streptomyces gramineus* | 1 |
|  |  | *Streptomyces griseorubens* | 3 |
|  |  | *Streptomyces koyangensis* | 1 |
|  |  | *Streptomyces longispororuber* | 4 |
|  |  | *Streptomyces maritimus* | 6 |
|  |  | *Streptomyces nigra* | 1 |
|  |  | *Streptomyces olivaceus* | 4 |
|  |  | *Streptomyces prasinosporus* | 1 |
|  |  | *Streptomyces pseudogriseolus* | 4 |
|  |  | *Streptomyces redgersensis* | 1 |
|  |  | *Streptomyces rochei* | 2 |
|  |  | *Streptomyces sampsonii* | 3 |
|  |  | *Streptomyces smyrnaeus* | 1 |
|  |  | *Streptomyces* sp. 102H11-4 | 1 |
|  |  | *Streptomyces* sp. 13650C | 8 |
|  |  | *Streptomyces* sp. CNS-753 | 1 |
|  |  | *Streptomyces* sp. FZ42 | 1 |
|  |  | *Streptomyces* sp. OAct 12 | 1 |
|  |  | *Streptomyces* sp. OAct 89 | 1 |
|  |  | *Streptomyces tempisqueensis* | 2 |
|  |  | *Streptomyces tendae* | 2 |
|  |  | *Streptomyces thermocarboxydus* | 1 |
|  |  | *Streptomyces variabilis* | 1 |
|  |  | *Streptomyces viridobrunneus* | 1 |
|  |  | *Streptomyces xylophagus* | 1 |
| *Nocardiopsaceae* | *Nocardiopsis* | *Nocardiopsis alba* | 21 |
|  |  | *Nocardiopsis dasonvelli* | 2 |
|  |  | *Nocardiopsis fildesensis* | 1 |
|  |  | *Nocardiopsis metallicus* | 3 |
|  |  | *Nocardiopsis salina* | 1 |
|  |  | *Nocardiopsis synnematoformis* | 3 |
| *Micromonosporaceae* | *Micromonospora* | *Micromonospora chalcea* | 2 |
|  |  | *Micromonospora maritima* | 1 |
|  |  | *Micromonospora* sp | 1 |
|  |  | *Micromonospora tulbaghia* | 1 |
| *Nocardiaceae* | *Rhodococcus* | *Rhodococcus aetherivorans* | 1 |
|  |  | *Rhodococcus cory* | 1 |
|  |  | *Rhodococcus rhodochrous* | 1 |
|  |  | *Rhodococcus zopfii* | 1 |
| *Pseudonocardiaceae* | *Actinomycetospora* | *Actinomycetospora chiangmaiensis* | 1 |
|  |  | *Actinomycetospora straminia* | 1 |
|  | *Pseudonocardia* | *Pseudonocardia kongjuensis* | 1 |
| *Micrococcaceae* | *Glutamicibacter* | *Arthrobacter mysoreus* | 1 |
|  | *Kocuria* | *Kocuria rosea* | 1 |
|  | *Micrococcus* | *Micrococcus aloeverae* | 1 |
|  | *Rothia* | *Rothia terrae* | 1 |
| *Brevibacteriaceae* | *Brevibacterium* | *Brevibacterium leteolum* | 2 |
| *Intrasporangiaceae* | *Kytococcus* | *Kytococcus sedenteris* | 1 |
| *Jonesiaceae* | *Jonesia* | *Jonesia denitrificans* | 1 |
| *Microbacteriaceae* | *Agrococcus* | *Agrococcus carbonis* | 1 |

From sponges, 53 actinobacterial isolates were obtained which belonged to 7 genera and 7 families. Whereas, 70 isolates were obtained from sediments that belonged to 7 different genera and 7 families. 17 isolates belonging to 8 genera of 7 families were isolated from sea water (Table 2). Even though, more number of isolates were obtained from sediment, Shannon’s diversity index showed higher putative species diversity in isolates obtained from sponge (Shannon’s H 3.27) than sediment (Shannon’s H 2.97) and water (Shannon’s H 2.76).

**Table 2:** Number of isolates belonging to different families obtained from various sources.

| **Families** | **Sediment** | **Water** | **Sponge** | **Total** |
| --- | --- | --- | --- | --- |
| *Actinomycetaceae* | 47 | 9 | 32 | **88** |
| *Brevibacteriaceae* | 1 | 1 | - | **2** |
| *Intrasporangiaceae* | 1 | - | - | **1** |
| *Microbacteriaceae* | - | 1 | - | **1** |
| *Micrococcaceae* | 1 | 1 | 2 | **4** |
| *Micromonosporaceae* | 1 | - | 4 | **5** |
| *Nocardiaceae* | 1 | 1 | 2 | **4** |
| *Nocardiopsaceae* | 18 | 2 | 11 | **31** |
| *Pseudonocardiaceae* | - | 2 | 1 | **3** |
| *Jonesiaceae* | - | - | 1 | **1** |
| **Grand total** | **70** | **17** | **53** | **140** |

More number of isolates (67) and more species diversity (10 genera with 43 putative species) were obtained from anthropologically less disturbed site Ade. Whereas, from Harnai which is a site with higher disturbance 47 isolates belonging to 25 putative species of 9 genera could be isolated. (Figure 2). Shannon’s diversity index (H) value for Ade was 3.45 and that for Harnai was 2.83. This indicates that the putative species diversity at Ade is more than diversity at Harnai.

### Production of Protease inhibitors

Results of PI production were recorded using spot assay as shown in Figure 3. It was observed that, PIs retained their activity even after heat treatment at 70°C for 15 mins. Out of 140 isolates used for screening of PIs, 113 isolates showed activity against at least one of the three pure proteases used for the study (viz. chymotrypsin, trypsin and subtilisin), whereas 27 isolates showed no PI production at all against any of these enzymes. Out of 113 isolates, 37 showed some degree of activity against all pure enzymes, of which 8 showed strong positive activity. Total of 17, 7 and 9 isolates showed PI activity exclusively against chymotrypsin, trypsin and subtilisin respectively (Figure 3). Majority of isolates which showed strong positive activity against enzyme chymotrypsin, also showed strong positive activity against subtilisin and trypsin. Further, even though 20% of the *Streptomyces* members showed complete absence of protease inhibitors, all the isolates showing strong positive activity against all enzymes belonged entirely to *Streptomyces* sp. Very few isolates of *Nocardiopsis* sp. showed PI activity against subtilisin. Regarding the source, highest number of strong positive isolates (7 out of 8) were obtained from the sediments and only 1 was obtained from the sponge. (Figure 2).

Out of 113 PI producers, strong positive (++) activity was shown by 10, 54, and 27 isolates against chymotrypsin, trypsin and subtilisin respectively (Table 3 A). A subset of 90 isolates belonging to most common genera *viz. Streptomyces sp*., *Nocardiopsis sp*. and *Micromonospora sp*. were chosen to check the activity against another extracellular bacterial protease which was obtained from cell free supernatant of *P. aeruginosa*, 36 isolates gave some degree of activity against this crude bacterial protease (Table 3B).

**Table 3A:** Genus-wise counts of PI producers active against pure proteases.

| Genus | Chymotrypsin |  |  | Trypsin |  |  | Subtilisin |  |  | Total |
| --- | --- | --- | --- | --- | --- | --- | --- | --- | --- | --- |
|  | - | + | ++ | - | + | ++ | - | + | ++ |  |
| *Actinomycetospora* | 0 | 2 | 0 | 1 | 1 |  | 2 | 0 | 0 | 2 |
| *Agrococcus* | 0 | 1 | 0 | 1 | 0 | 0 | 1 | 0 | 0 | 1 |
| *Brevibacterium* | 1 | 1 | 0 | 1 | 1 | 0 | 2 | 0 | 0 | 2 |
| *Glutamicibacter* | 0 | 1 | 0 | 0 | 0 | 1 | 1 | 0 | 0 | 1 |
| *Jonesia* | 0 | 1 | 0 | 1 | 0 | 0 | 1 | 0 | 0 | 1 |
| *Kocuria* | 0 | 1 | 0 | 1 | 0 | 0 | 1 | 0 | 0 | **1** |
| *Kytococcus* | 0 | 1 | 0 | 0 | 1 | 0 | 1 | 0 | 0 | **1** |
| *Micrococcus* | 0 | 1 | 0 | 0 | 1 | 0 | 1 | 0 | 0 | **1** |
| *Micromonospora* | 2 | 3 | 0 | 2 | 1 | 2 | 4 | 1 | 0 | **5** |
| *Nocardioopsis* | 6 | 23 | 2 | 17 | 3 | 11 | 27 | 4 | 0 | **31** |
| *Pseudonocardia* | 0 | 1 | 0 | 1 | 0 | 0 | 1 | 0 | 0 | **1** |
| *Rhodococcus* | 0 | 4 | 0 | 2 | 1 | 1 | 4 | 0 | 0 | **4** |
| *Rothia* | 0 | 1 | 0 | 0 | 0 | 1 | 1 | 0 | 0 | **1** |
| *Streptomyces* | 43 | 37 | 8 | 31 | 19 | 38 | 33 | 28 | 27 | **88** |
| **Total** | **52** | **78** | **10** | **58** | **28** | **54** | **80** | **33** | **27** | **140** |

**Table 3B:** Genus-wise counts of PI producers active against crude protease.

| Genus | Crude *P. aeruginosa* Protease |  |  | Total |
| --- | --- | --- | --- | --- |
|  | - | + | ++ |  |
| *Micromonospora* | 2 | 1 | 0 | 3 |
| *Nocardioopsis* | 3 | 4 | 0 | 7 |
| *Streptomyces* | 49 | 29 | 2 | 80 |
| **Grand Total** | **54** | **34** | **2** | **90** |

### Effect of Season and Anthropological Activities on Production of Protease Inhibitors

Least number of actinobacterial isolates were obtained from monsoon and post-monsoon seasons and the number of isolates obtained from each sampling increased towards the month of May i.e. the pre-monsoon season. Majority of actinobacterial isolates were obtained from pre-monsoon season (Figure 4A). From Ade and Harnai, over all, occurrence of inhibitors against chymotrypsin and trypsin was more as compared to that against subtilisin (Figure 4B). Proportion of isolates not giving PI activity against any of the enzymes was higher in Ade (14 out of 67 isolates i.e. approximately 21%) compared to that in Harnai (4 isolates out of 47 i.e. approximately 19%) (Figure 4C). Thus to summarize, more number of putative species were obtained from Ade however, proportion of PI producers from that site was less compared to Harnai.

**Fig 4:**
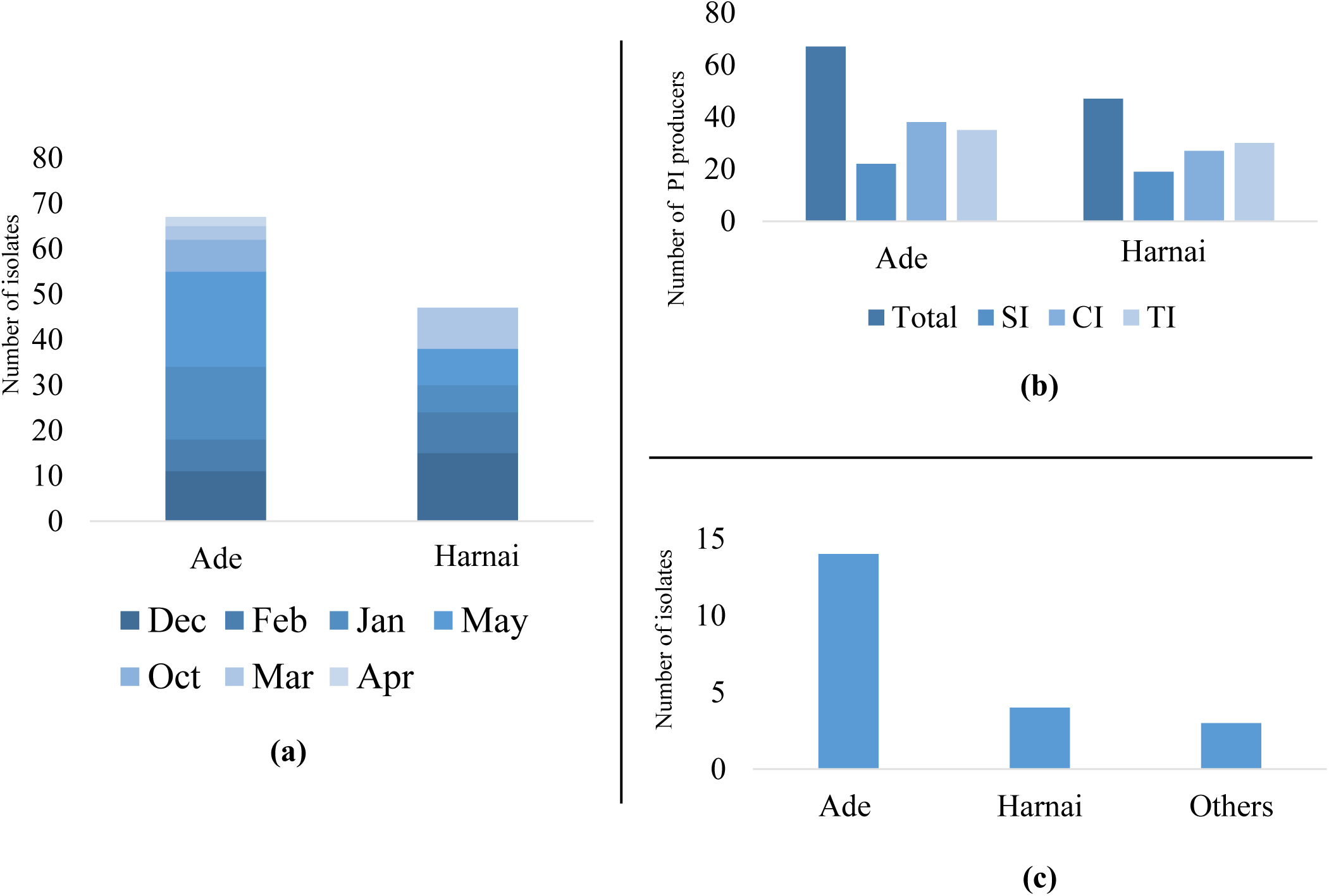
Graphs showing number of actinobacterial isolates obtained from Ade and Harnai in different seasons (A), number of isolates showing inhibitory activity against subtilisin (SI), chymotrypsin (CI) and trypsin (TI) is shown in (B) and number of isolates negative for PI production are shown in (C).

## DISCUSSION

Actinobacteria are widespread in terrestrial and aquatic environments. Many of them also produce external spores which are resistant to dehydration (Trujillo, 2016). Various reports say that marine actinobacteria have a high potential for production of biomolecules (Jose and Jha, 2017; Solanki et al., 2008) however, the number of actinobacterial compounds discovered so far are limited not by the number of compounds produced by them but by the amount of screening efforts put in (Watve et al., 2001). Many reports have highlighted the need for studying marine actinomycete diversity and their potential use for obtaining novel metabolites (Dharmaraj, 2010; Jensen et al., 2005; Lam, 2006; Subramani and Aalbersberg, 2012). Researchers from India have also isolated actinobacteria from coastal regions of West Bengal (Peela et al., 2005; Ramesh and Mathivanan, 2009), Gujarat (Jose and Jha, 2017), Tamil Nadu (Raja et al., 2010; Valli et al., 2012) and Andaman and Nicobar (Karthik et al., 2014) to name a few for bioprospecting studies. However, there are very few reports of isolation of actinobacteria and still fewer reports on PI producing actinobacteria from the coast of Maharashtra. Thus considering these facts, current study focused on isolation of actinobacteria from coast of Maharashtra and their potential as producers of PIs.

### Phylogenetics of Actinobacteria from Intertidal Regions of Maharashtra

From 7 different sites including Ade and Harnai used in the study, high diversity of actinobacterial species was obtained. Highest number of isolates were obtained from sediment samples followed by sponge and water samples. Majority of isolates belonged to *Streptomyces sp*. followed by *Nocardiopsis sp*. and *Micromonispora sp*. The observations coincided with observations of (Jose and Jha, 2017) who worked on the samples obtained from coast of Gujarat which is situated north of Maharashtra. These findings might suggest that members of above three genera have adapted well to live in marine environments and show a widespread distribution along the western coast of India. Few earlier reports strongly support the existence of sponge‐specific microbes (Simister et al., 2012; Taylor et al., 2007). In the current study, higher Shannon index values for sponge associated actinobacteria likely suggests their sponge specific nature.

### PI Production by Marine Actinobacteria

Marine actinobacteria are known to produce large number of bioactive molecules like antimicrobial and anticancer compounds intra-as well as extracellularly (Subramani and Aalbersberg, 2012), however, it is difficult to understand the strategy of producing extracellular molecules in the marine ecosystems, since, the molecules secreted outside can be easily diluted in the surroundings. Endopeptidases like chymotrypsin and trypsin are also secreted extracellularly by bacteria and invertebrates in the marine environments (Holmström and Kjelleberg, 1999; Thao et al., 2015). Moreover, reports have shown that coastal waters contain significant amounts of trypsin-type and chymotrypsin-type endopeptidases (Obayashi and Suzuki, 2005). Interestingly, results of current study demonstrated that high number of actinobacteria produced PIs against chymotrypsin and trypsin. Further, more than 80% of cultured actinobacteria produced extracellular inhibitors against at least one of the four enzymes and almost 20% isolates showed inhibition against all the enzymes used in the study.

The presence of PIs suggests that they are needed for the defence of organisms against exo- and endoproteases present in the surrounding environment. Moreover, they might be used as defence molecules against proteases secreted by protists and bacteria.

### Micro-ecological Dynamics in PI Producing Actinobacteria

In this study, PI producing actinobacteria were obtained from all locations and from samples collected in all seasons. However, there was a remarkable difference in proportions of PI producing actinobacteria obtained from sites with varying human disturbance. As seen from Shannon’s diversity index, comparatively less disturbed site Ade showed more species diversity, majority of non-producers of PI were reported from this site. On the contrary, almost 80% isolates obtained from Harnai (which has more human interference and thus chances of containing higher organic load) were producing PIs. Organic load and presence of extracellular proteases in the surroundings might be a driving force for production of extracellular PIs. Therefore, these results can be used as a lead for carrying out studies on a larger expanse to check correlation of organic load and production of PIs by the marine actinobacteria.

Actinobacteria are involved in bacterial predation at oligophilic conditions (Kumbhar et al., 2014), they might need to produce proteases for killing prey species. Hence, one of the possibilities of finding high number of PI producers amongst actinobacterial isolates might be to achieve self-protection so that their own proteases are rendered ineffective against their own selves. Hence, it would be interesting to check if there is any significant correlation between production of PI and bacterial predation, (Baig et al., 2020).

Through this study we show that extracellular protease inhibitor producing actinobacteria are abundant in the intertidal zones of west coast of Maharashtra. We hypothesize that this might have a correlation with extracellular proteases and bacterial load in marine environment. Understanding the need for production of protease inhibitors by actinomycetes can give interesting insights into marine microbial ecology and clues for production of novel protease inhibitors.

## ACKNOWLEDGEMENT

Financial support for this project was provided by Rajiv Gandhi Science and Technology Commission under the Maharashtra Gene Bank Program. We thank Dr. Milind Watve and Dr. Neelesh Dahanukar for their critical comments and suggestions for this manuscript. We also thank Mr. Asim Auti for providing *Pseudomonas aeruginosa* culture and Mr. Chinmay Kulkarni for assisting in the fieldwork.

